# Platelet phosphorylated TDP-43: An exploratory study for a peripheral surrogate biomarker development for Alzheimer’s disease

**DOI:** 10.1101/132837

**Authors:** Rodger Wilhite, Jessica Sage, Abdurrahman Bouzid, Tyler Primavera, Abdulbaki Agbas

## Abstract

Aim: Alzheimer’s disease (AD) and other forms of dementia create a non-curable disease population in World’s societies. To develop a blood-based biomarker is important so that the remedial or disease-altering therapeutic intervention for AD patients would be available at the early stage. Materials & Methods: TDP-43 levels were analyzed in post-mortem brain tissue and platelets of AD and control subjects. Results: We observed an increased TDP-43 (<60%) in post-mortem AD brain regions and similar trends were also observed in patient’s platelets. Conclusion: Platelet TDP-43 could be used as a surrogate biomarker that is measurable, reproducible, and sensitive for screening the patients with some early clinical signs of AD and can be used to monitor disease prognosis.

**Lay abstract:** In this study, we explore to identify an Alzheimer’s disease-selective phospho-specific antibody that recognizes the diseased form of TDP-43 protein in patient’s blood-derived platelets. Our results suggest that selective anti-phosphorylated TDP-43 antibody discriminates Alzheimer’s disease from non-demented controls and patients with amyotrophic lateral sclerosis. Therefore, platelet screening with a selective antibody could potentially be a useful tool for diagnostic purposes for Alzheimer’s disease.

## INTRODUCTION

More than 5 million Americans are living with Alzheimer’s disease (AD) and this number is projected to rise to 13.8 million by 2050. As the world population gets older, the incidence of AD is rising. Unfortunately, only 1 in 4 people with AD have been diagnosed [1].The rising cost of health care for AD patients has a negative socioeconomic impact on global society as well as being a burden on caretakers. The early diagnosis of AD could be critical for starting an effective treatment with current options as well as designing new competent disease-modifying approaches. There is a great need to improve early detection in the course of neurodegenerative diseases such as AD, Amyotrophic lateral sclerosis (ALS), Parkinson’s disease (PD), frontotemporal lobar disease (FTLD), and others so that the timely application of disease-specific treatments would be effective. Current diagnostic tests for AD rely on expensive brain imaging technology that is available only to a few patients [2]; cognitive and psychiatric assessments; the collection of cerebrospinal fluid (CSF) samples, which requires invasive procedures; and lumbar puncture that has a negative public perception in several countries [3]. Therefore, it becomes an urgent task to identify better biomarkers for AD as well as for other neurodegenerative diseases. More specifically, new biomarkers should be sensitive, selective, reliable, affordable, and involve a non-invasive sampling method. Such biomarkers are in great demand for the early stages of neurodegenerative diagnosis and making dementia screening a viable approach. We have turned to explore for a biomarker molecule(s) that can be measurable in blood tissue. There are several fluid-based biomarker candidates for AD [4-7]; however, either the milieu of the diagnostic biomolecules or the measurement platforms for them have made many of these biomarkers unfavorable candidates. Furthermore, the popular AD biomarker candidates plasma Aβ oligomers and *tau* were recently found to be not good contenders due to measuring environment [8, 9]. Serum/plasma contains substantial amount of albumin and immunoglobulins, which interfere with the measurement of the target protein. Removal of interfering biomolecules could easily reduce the measuring of AD-relevant signature proteins. These reported observations led us to explore new blood-based surrogate biomarker(s) that may be useful for AD diagnosis.

One new potential biomarker candidate for AD is **T**rans-**a**ctivation **r**esponse DNA/RNA binding protein (TARDP). Due to its 43 kDa size, the acronym TDP-43 will be used throughout this paper. Substantial research has been conducted to decipher the role of TDP-43 in different cellular events as well as its role(s) in neurodegenerative disease states [10-14]. TDP-43 is ubiquitously expressed in all nucleated cells [15, 16], and it has the ability to shuttle in-and-out between nucleus and cytoplasm due to having nuclear localization and nuclear export sequences [17-20]. Cytosolic TDP-43 is not well described yet; however, cytosolic phosphorylated derivatives of TDP-43 in neurodegenerative diseases could be responsible for hyperphosphorylated aggregate formation and could be considered as a biomarker candidate. C-terminus of TDP-43 is much enriched with Ser amino acids which are potential targets for phosphorylation as predicted by PONDR ® analysis (Fig. 1A). In neurodegenerative diseases such as AD, FTDL, and ALS, the tissue levels of TDP-43 are increased, mostly located in the cytosol, and consistently observed in inclusion bodies in neurons [16] rather than the nucleus. In Pick disease (PiD), the presence of TDP-43 inclusions suggests that TDP-43 accumulation and modification are an important component of PiD [21]. Post-translationally modified TDP-43 aggregates were also observed in post-mortem brain tissue sections [15]. TDP-43 positive inclusion bodies are becoming more detectable, seen in about 75% of the brain tissues from AD patients [11, 22]. The elevated level of TDP-43 in blood suggests that TDP-43 may be considered as a potential surrogate biomarker in neurodegenerative diseases [23, 24]. TDP-43 is prone to phosphorylation and undergoes truncation if it remains in the cytosol [25]. A recent study has provided some evidence that TDP-43 mislocalization was an early or pre-symptomatic event and was later associated with neuron loss [26]. These studies suggest that post-translationally modified (i.e. phosphorylated) TDP-43 could be viewed as a disease-specific protein. Therefore, we have focused on measuring phosphorylated derivatives of TDP-43 in platelets, because it represents the diseased form of cytosolic TDP-43.

**Fig. 1A.**
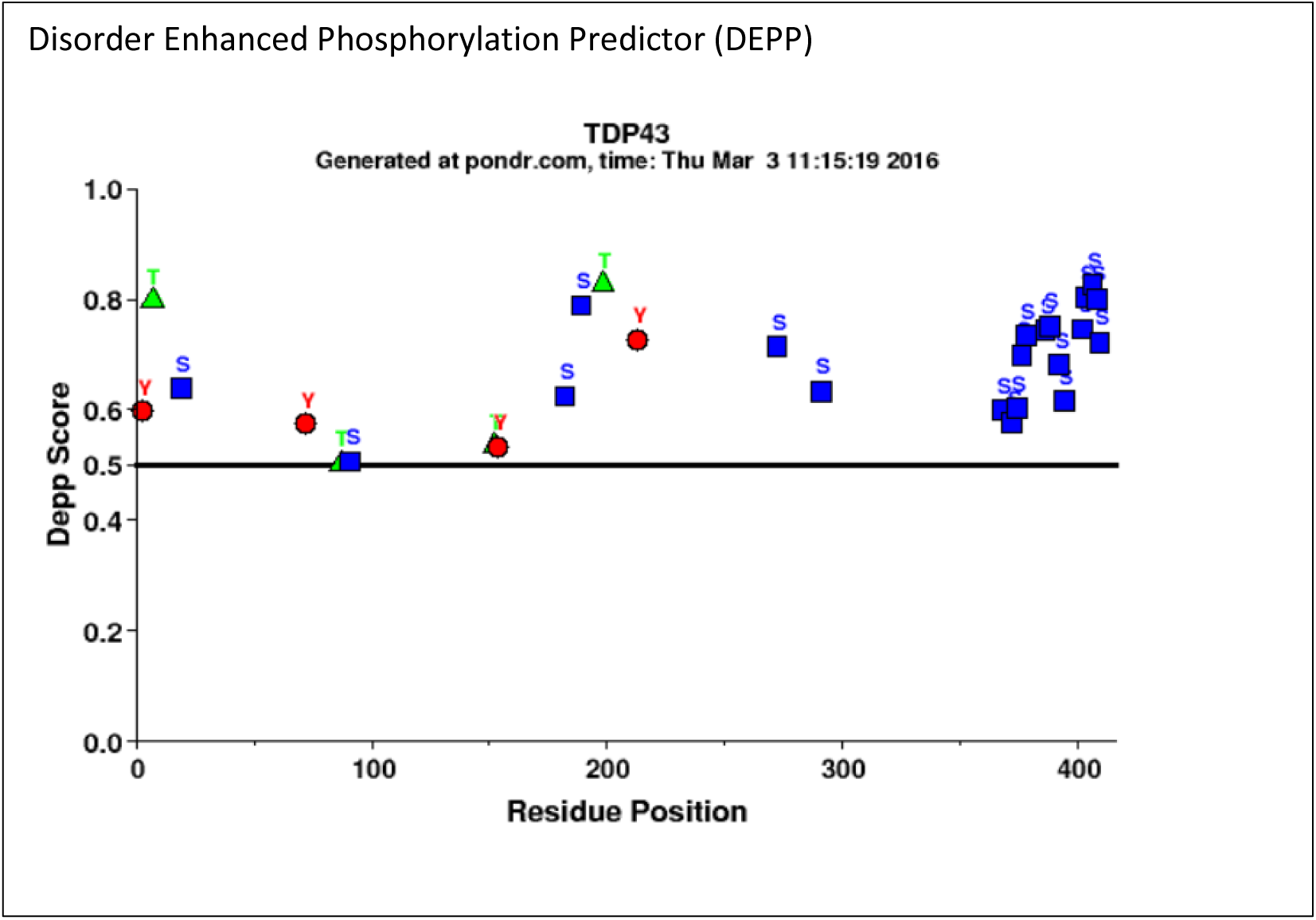
PONDR® analysis of TDP43 for potential phosphorylation sites. Majority of the phosphorylation events were predicted at Serine (Ser) amino acid sites (359-410). Most of the Ser amino acids are located at C-terminus region (20 out of 41; 48.7%) *www.molecularkinetics.com;*) under license from the WSU Research Foundation. PONDR® is copyright (c)1999 by the WSU Research Foundation, all rights reserved.

**Fig.1B.**
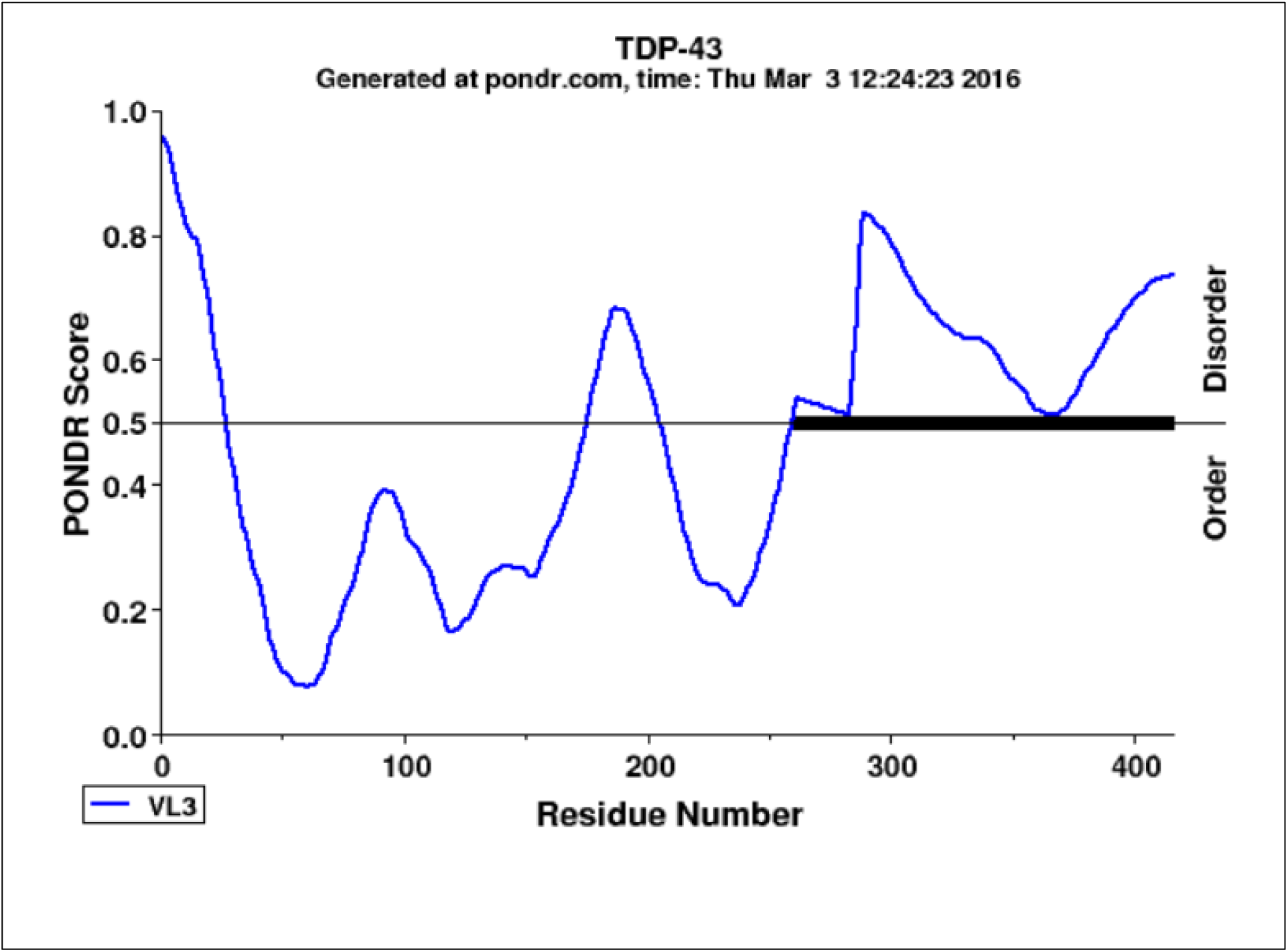
PONDR® VL3-BA analysis of TDP43 for identifying the disordered sites. The VL3-BA predictor is a feedforward neural network that was trained on regions of 152 long regions of disorder that were characterized by various methods. The region close to C-terminus was identified as disordered sites for TDP-43 which is also most of the Serine amino acids are located. *www.molecularkinetics.com;*) under license from the WSU Research Foundation. PONDR® is copyright (c)1999 by the WSU Research Foundation, all rights reserved.

Monitoring disease-relevant biomolecules in the central nervous system (CNS) is a challenge due to unfeasibility and invasiveness of taking repeated samples from brain and spinal cord tissues. Hence, sampling blood platelets may serve as a feasible media where the aberrant TDP-43 is detectable. Platelets are anuclear blood cell fragments that are derived from megakaryocytes [27] and share the biochemical properties of neurons [28-30]. Accordingly, several investigators consider using platelets as a venue to study the pathogenesis of neurodegeneration. The latest progress on platelets as a reliable source for biomarkers was reviewed in the literature [29, 31-35]. One of the notable items of research was conducted on platelet-derived secreted amyloid precursor protein-β (APP-β) for AD [35]; however, their data does not tell much about which Aβ peptide was measured.

## MATERIAL METHODS

### Reagents

Anti hTARDBP polyclonal antibody (ProteinTech Group, Chicago, IL; Cat#1078-2-AP) and phosphorylated derivatives of the pTDP-43 antibodies (CosmoBio USA; Cat#TIP-TD-P09, TIP-TD-P07, TIP-PTD-P05, TIP-PTD-P03, TIP-PTD-M01, TIP-PTD-P01, TIP-PTD-P02, TIP-PTD-P04) (Abcam Cat# Ab184683, ProteinTech Cat# 10782-2-AP,66318-1-Ig,22309-1-p(discontinued); Sigma Cat# T1705,SAB4200225; Biolegend Cat# 829901) were commercially purchased. Citrate Wash Buffer (11mM glucose, We chose platelets for identifying and measuring both total and phosphorylated TDP-43 protein derivatives in AD for the following reasons: (i) platelets are easy to repeatedly obtain from the patients with minimal distress; (ii) their life span is short (7-10 days) [36] which will reflect dynamic changes on phosphorylated TDP-43; (iii) it was reported that platelets transiently open the blood brain barrier (BBB) [37]. Consequently, biomolecules may come in to contact with the blood stream and become absorbed by platelets; and (iv) serum/plasma proteins and other biomolecules are exposed to dilutions and result in large analytical challenges; platelet content is protected from such changes. We have hypothesized that platelet TDP-43 and its phosphorylated derivatives may be considered as a viable surrogate dynamic biomarker. The elevated brain TDP-43 protein species in AD patients can be determined in the platelet lysate so that platelet TDP-43 readings could be used as a surrogate biomarker for AD that helps to monitor the progress of disease during the treatment.

128mM NaCl, 4.3 mM NAH_2_PO_4_, 7.5 mM Na_2_HPO_4_, 4.5 mM sodium citrate, and 2.4mM citric acid, pH 6.5)[38] and platelet rupture Buffer (250 mM sucrose, 1 mM EDTA, 10 mM Tris, pH 7.4) were prepared in our lab using reagent grade chemicals. Phosphatase inhibitor cocktail (Calbiochem # D00147804) (1:1,000) and protease inhibitor cocktail (Calbiochem# 539134) (1:2,000) were added to the platelet rupture buffer just before use to preserve TDP-43 proteins from proteolytic degradation and dephosphorylation processes.

### Sample preparations

#### Human Platelets

Human blood-platelet samples were obtained from the following sources; (1) The Bio-specimen Bank of University of Kansas Medical Center (KUMC); platelets were previously collected from AD patients and age-matched with otherwise healthy subjects (non-demented) and stored at -80°C and (2) ALS clinic at the University of Kansas Medical Center, Kansas City. ALS patient platelets were utilized as a negative disease control for identifying a selective antibody for AD patient platelets. AD patients in this study were not classified early onset (EOAD) vs. late onset (LOAD). The genotyping for Amyloid precursor protein (APP) and Presenilin (PSEN) associated EOAD was not performed. All AD patients (n=10) were categorized based on age-relevant cognitive symptom onset. The baseline evaluation included a thorough clinical examination by a trained clinician was conducted. This evaluation included a Clinical Dementia Rating (CDR) to exclude the presence of dementia was performed according to clinical and cognitive variables and descriptive data collected from Alzheimer Disease Centers [39]. A trained psychometrician administered a comprehensive (National Alzheimer’s Coordinating Center Uniform Data Set, versions 2 and 3) cognitive testing battery [39]. Clinical and psychometric test results were reviewed and discussed at a weekly consensus conference that included clinicians, a neuropsychologist, and raters. The ALS patients were clinically diagnosed by physicians and the subject identities for the biosamples were de-identified. All patients and otherwise healthy individuals were given a consent form before obtaining the blood samples. The sample collection procedure was approved by the Institutional Review Board of Kansas City University of Medicine and Biosciences (KCU) and the University of Kansas Medical Center (KUMC).

Platelet donors comprised five female AD patients with average age of 75.2±10.5 and five male AD patients with average age of 75.8 ±8.6. Non-demented control subjects comprised two males with average age of 77.5 ±2.13 and eight non-demented females with average age of 73.6 ± 3.5.

Platelets were isolated from freshly drawn blood into ACD containing vacutainer tube from clinically diagnosed patients and otherwise healthy subjects according to a standard two-step low speed centrifugation method adopted by Bio-Specimen bank facilities of the University of Kansas Medical Center. The platelet pellets were gently washed with 1 ml of citrate was buffer [38] and re-centrifuged at 200xg for 20 min. The washed platelet pellet was ruptured, sonicated in 0.6 ml of rupturing buffer with protease and phosphatase inhibitors, and subjected to high speed centrifugation (16,000 x g; 30 min; 4°C) to obtain platelet cytosol. Protein concentrations were determined by the bicinchoninic acid (BCA) spectrophotometric method [40]. The samples were aliquot and stored at -80°C until use.

#### Human Brain Sample preparation

The human brain tissue samples from post-mortem AD patients and age-matched control subjects were obtained from the Bio-Specimen bank of the University of Kansas Medical Center. The 100-200 mg samples of excised tissue from three brain regions (frontal cortex, cerebellum, and hippocampus) were received and homogenized in a Teflon-pestle glass homogenizer containing. ice-cold buffer (0.32 M sucrose, 0.5 mM MgSO_4_, 10 mM epsilon-caproic acid, 0.1 mM EGTA, protease inhibitor cocktail 0.1% v/v, 10 mM HEPES, pH 7.4). Tissue : Homogenate buffer ratio was kept at 1:20 for achieving efficient solubilization The homogenization was carried out in an ice bucket with 8-10 strokes. The homogenate was aliquoted and stored in -80°C until use. Protein concentrations were analyzed by the BCA method [40].

#### Western Blot

The brain homogenate and platelet lysate proteins (20-30 µg/well) were resolved in 12 % SDS-PAGE and 4-20% SDS-PAGE, respectively under the reducing conditions. The proteins were transferred onto a PVDF membrane and subsequently the membrane was probed with both pan anti-TDP-43 and several anti-phosphorylated TDP-43 antibodies listed in reagents section. The protein bands were visualized by enhanced chemiluminescence and infrared dye based fluorescence methods, and they were analyzed by NIH’s ImageJ (V.1.46r) and Image Studio^TM^ Lite software (V. 4.0)

#### Capillary Electrophoresis

The platelet lysates from AD, ALS patients and otherwise healthy subject cohort were analyzed by a simple western system, a new technology developed by ProteinSimple, Inc., USA. This technology does not require classical SDS/PAGE and Western blotting components. It uses very little sample-mix volume (∼3-5 µl) and provides high-throughput style sample run. The platelet lysate proteins (0.2 mg/ml) were analyzed in duplicate and both capillary electropherogram and pseudo protein bands were generated and analyzed by the system software (Compass for Simple Western, v.3.0.9).

#### Statistical analysis

Mann-Whitney U rank sum test was employed for statistical analysis.

## RESULTS

### TDP-43 protein level was differentially increased in AD-patient brain tissue and this trend was reflected in platelets

In early stages of this work, we have shown that total TDP-43 protein levels were increased in the brain regions of post-mortem AD patients (n=3). The most noticeable TDP-43 increase was observed in the hippocampus while the frontal cortex and cerebellum reflected a slight TDP-43 increase as compare to non-symptomatic control subjects (Fig. 2A). Total TDP-43 aggregates were observed in three different brain regions and the most notable aggregates were observed in the hippocampus (Fig.2B). We have also noticed that the platelet lysate TDP-43 levels were increased by <60 % in AD patients (n=3) (Fig. 2C) in the early phase of this study. Readers should be advised that platelet lysates were obtained from a separate AD patient cohort, because the University of Kansas Medical Center Bio-specimen repository did not have the matching post-mortem tissue and platelet lysates from the same AD patients and non-symptomatic control individuals.

**Fig. 2A.**
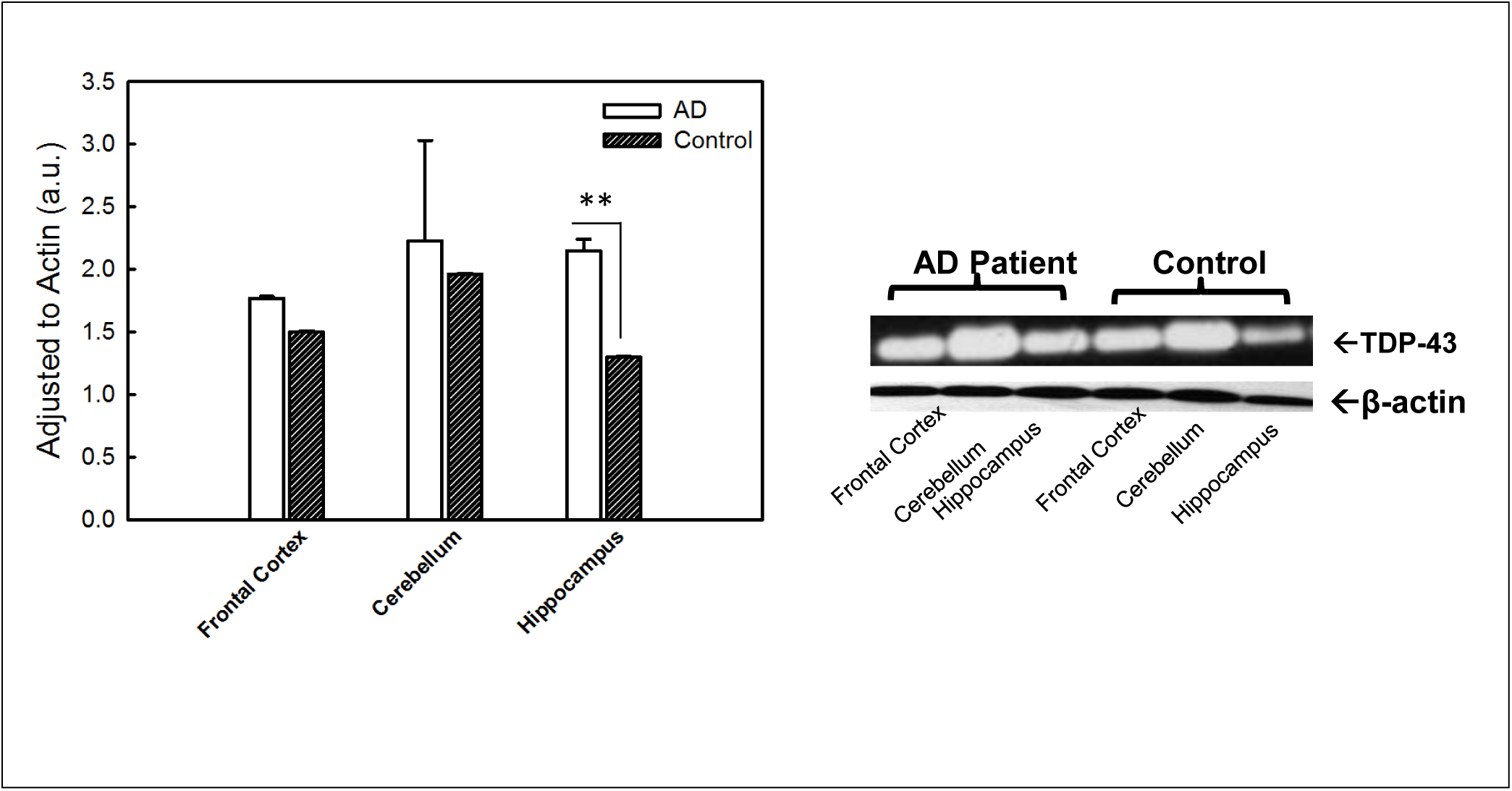
TDP-43 distribution in Alzheimer’s disease (AD) patient brain regions. The tissue from three different regions of the brain was used in this study. The tissue homogenates were analyzed by immunoblotting method. The protein band intensities were normalized to actin. Three post-mortem AD patients and age-matched healthy human brain samples were utilized in this study (n=3). The difference between control and AD in hippocampus region was found statistically significant (P≤0.015) according to Mann-Whitney U rank sum test.

**Fig.2B.**
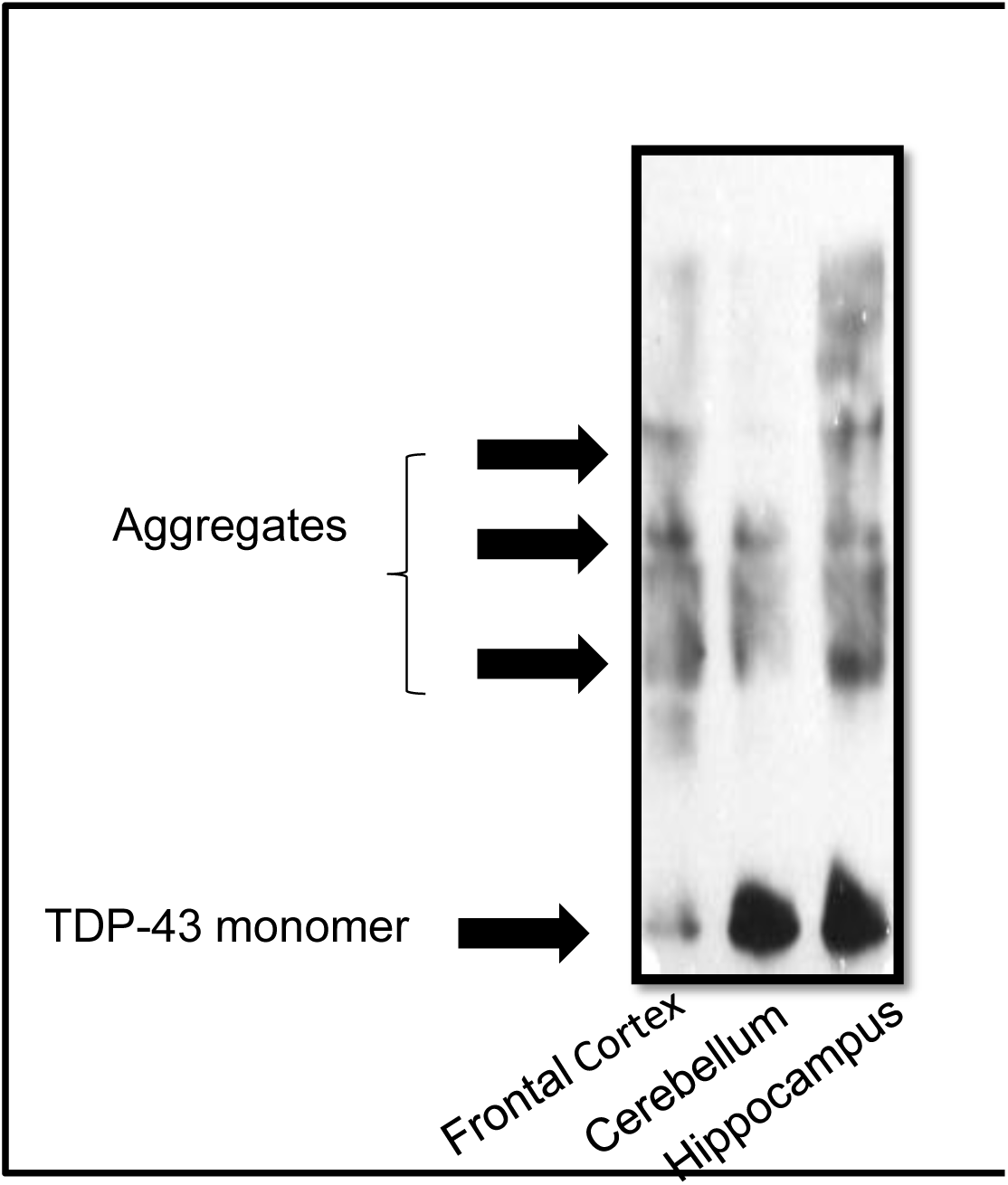
TDP-43 protein aggregation in Alzheimer’s disease patient’s brain region. The homogenates from different regions of the brain were resolved in non-reducing SDS/PAGE condition and immunoprobed with anti-TDP-43 (pan) antibody (1:1000 dilutions). The TDP-43 protein aggregation is relatively more prominent in hippocampus region.

**Fig. 2C.**
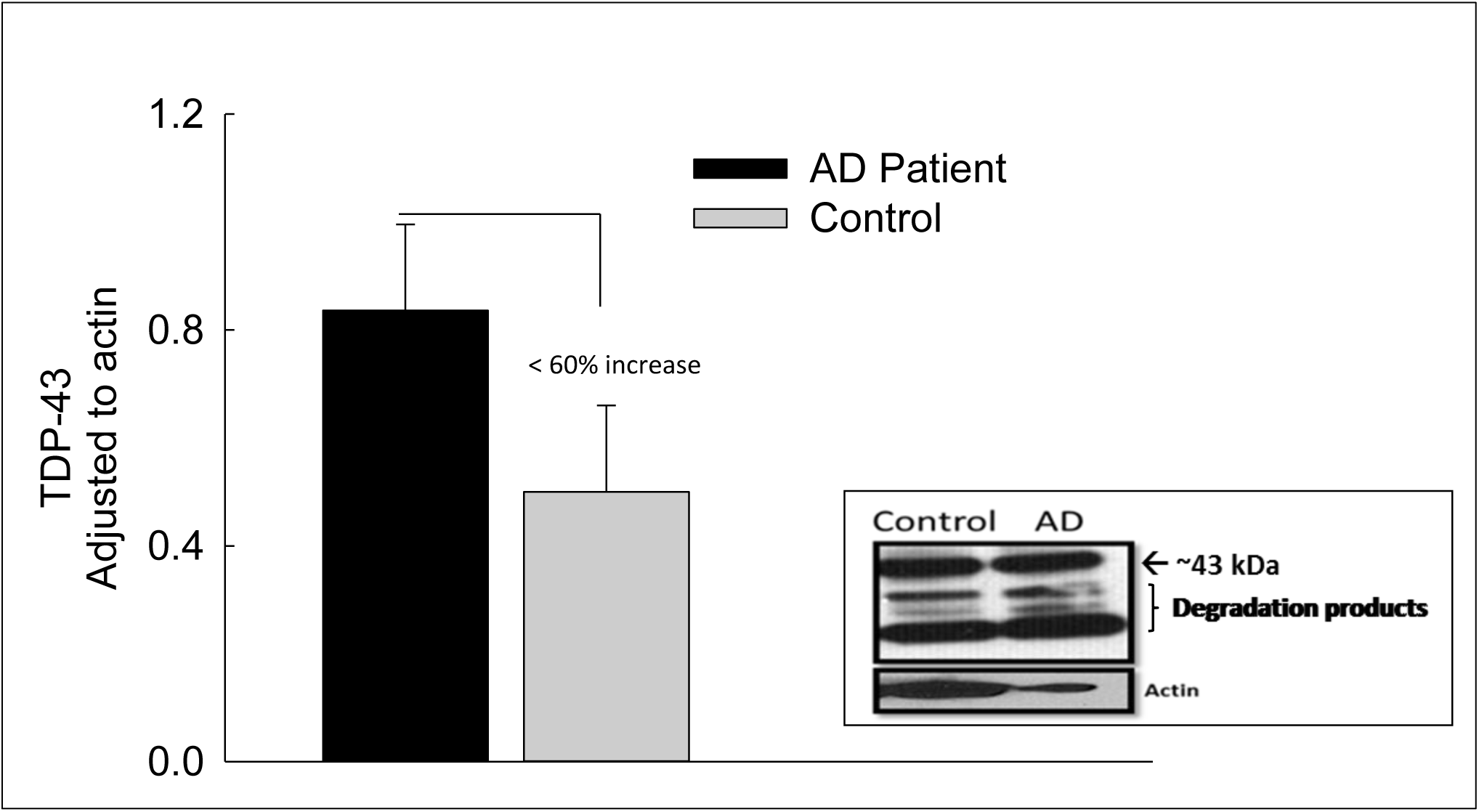
Platelet lysate TDP-43 profile. This figure represents the early TDP-43 studies on platelets obtained from AD patients (n=3) and age-matched healthy subjects (n=3). The platelet lysates were analyzed by a classical immunoblotting assay using anti-TDP-43 (pan antibody (1:1000 dilutions).

### A sequence-specific anti-phosphorylated TDP-43 Ab distinguished AD from other neurodegenerative disease

In the next phase of this study, we have focused on identifying an AD selective anti-phosphorylated TDP-43 Ab as a screening tool in a small number of subject platelets (n= 10 in each group). First, we have employed a computer based **P**redictor **o**f **N**atural **D**isordered **R**egion (PONDR®) algorithm using TDP-43 sequence (NCBI accession code: Q5R5W2.1). Disordered Enhanced Phosphorylation Predictor (DEPP) analysis predicted 28 potential phosphorylation sites and a majority of them were Ser amino acid enriched on the C-terminus (aa 369-410) (Fig.1A). Another algorithm (PONDR® VL3-BA) was employed to predict 152 aa long regions of disorder that were characterized by other methods (Fig.1B). Nuclear magnetic resonance (NMR) studies also revealed that a ∼ 80 aa sequence from the C-terminus region of TDP-43 was identified as the most disorderly region [41, 42] where the majority of phosphorylation sites were located. Therefore, we have tested several anti-phosphorylated TDP-43 antibodies, raised for several different sequence of the C-terminal of TDP-43 protein, purchased from various vendors (ProteinTech, Abcam, Cosmobio-USA, Sigma, and Biolegend) to identify an AD-selective antibody that can be used for screening assays. An anti-phospho (S409/410-2) TDP-43 antibody (ProteinTech Cat# 22309-1-AP) was identified as a potential antibody that discriminates AD platelet lysate phospho-TDP-43 profile from that of amyotrophic lateral sclerosis (ALS) (negative disease control) (Fig. 3A) and from that of non-symptomatic, otherwise healthy age-matched subjects (Fig.3B). A prominent protein peak at about 62 kDa position was consistently observed in AD patient platelet lysates (Fig.3A).

**Fig. 3A.**
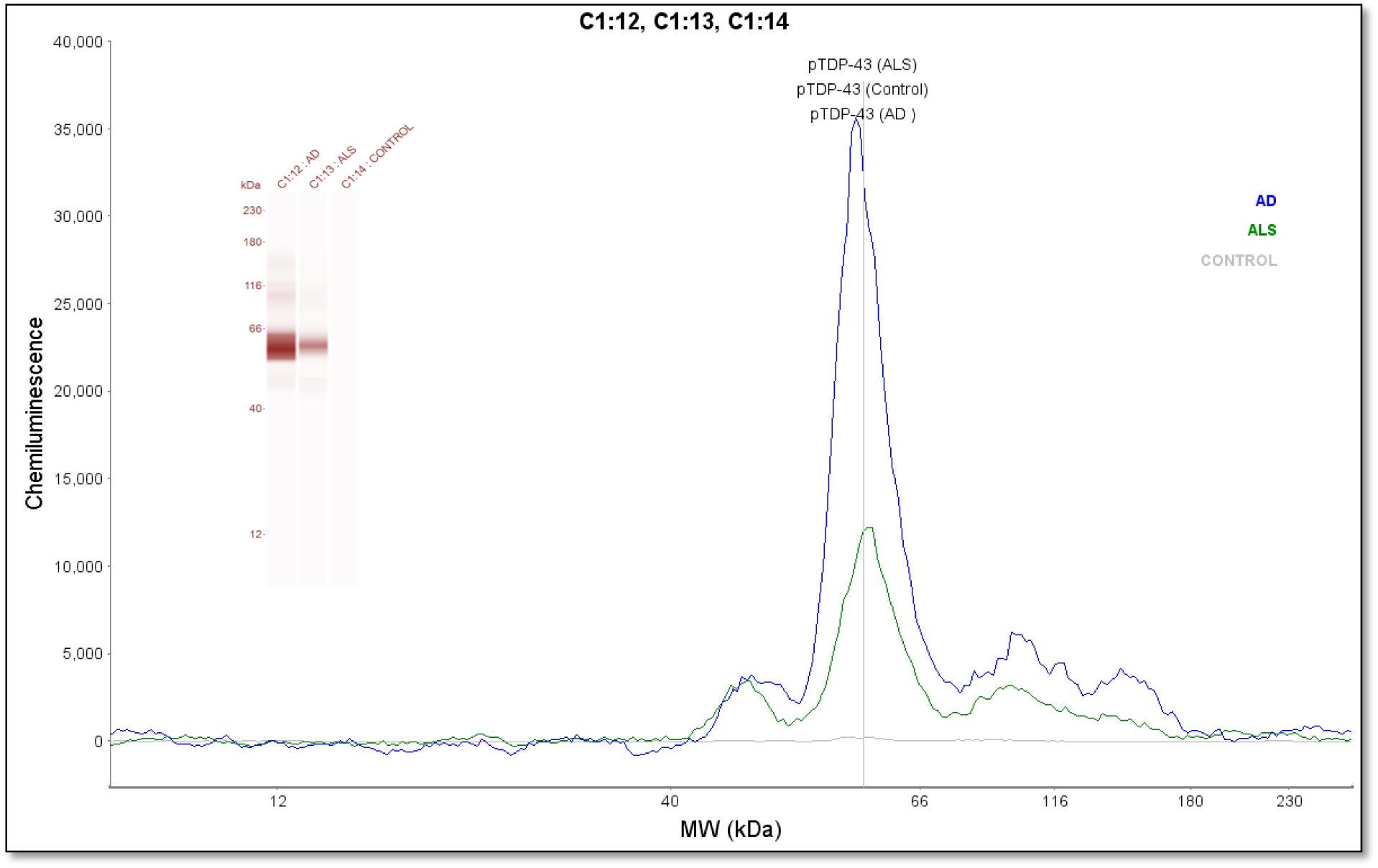
Phosphorylated TDP-43 profile in platelet lysates. An anti-phospho (Ser409/410-2) TDP-43 antibody was used as an immunoprobing agent. The signals from AD platelet lysates was more pronounced as compare to ALS (negative disease control) and healthy subjects (control). Inset figure shows a computer generated pseudo band that marks prominent phosphorylated TDP-43 at about 62 kDa.

**Fig. 3B.**
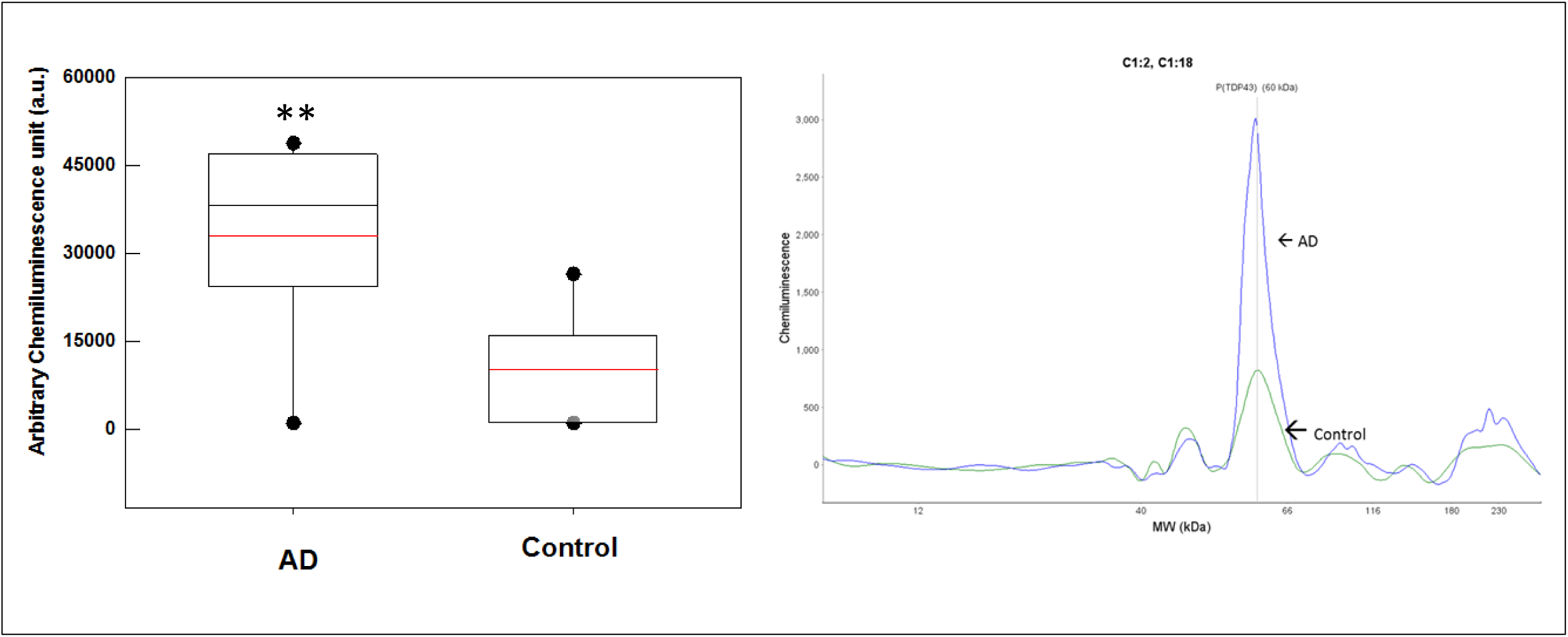
Platelet lysate phosphorylated TDP-43 protein profile. The platelet lysates obtained from local biorepository (n=10 in each group) and analyzed by capillary electrophoresis based gel-less and membrane-less system western assay developed by Proteinsimple (WES). The electropherogram peaks indicate the noticeable difference at about 62 kDa protein that represents phosphorylated TDP-43 protein. Box-whiskers plot represents statistical values. Redline within the boxes mark the median. Mann-Whitney U rank sum test was employed for statistical analysis. Difference between AD and control platelet phosphorylated TDP-43 was found statistically significant (P≤ 0.010)

## DISCUSSION

Misfolded aberrant protein aggregation is frequently observed in neurodegenerative diseases [43-45]. Pathologically misfolded protein aggregate formation occurs long before any measurable cognitive decline [46-48]. Current brain imaging methods are expensive and not easily accessible in many medical facilities. Therefore, it is essential to develop a blood-based biomarker assay platform that is feasible, cost-effective, and selective. This assay may aid medical evaluations to predict AD before the clinical manifestations are revealed as well as to monitor the potential AD treatments.

In this preliminary study, we have tested our hypothesis and provided some new findings that human blood-derived platelet TDP-43 and its derivatives may reflect the changes in the TDP-43 profile in human AD brain. There are several reports in the literature supporting the link between TDP-43 pathology and diagnosed AD cases [11, 49-51]. In early stage of this study, we have demonstrated TDP-43 protein aggregation in select brain regions (Fig.2A, **2B**). Hippocampus had relatively high level of TDP-43 aggregation as compare to the other brain region, and the difference was found statistically significant (P≤0.015; Mann-Whitney U rank sum test) (Fig. 2A). We are aware of the insufficient sample numbers (n=3) analyzed in this study. It was a challenge for us to obtain more AD patient brain samples. We continue to communicate with bio-repositories in other health centers to obtain more brain tissues. We have shown the hippocampus region was very vulnerable to oxidative stress in the aging process [52]. This also partially explains why increased level of TDP-43 aggregation was observed in the hippocampus region, which is the most important region for memory storage.

In the next stage of this study, we tested whether AD patient brain TDP-43 protein profile could be observed in peripheral tissues such as blood-derived platelets. If we could demonstrate the same or similar profile in platelets, then we will have an assay platform by which TDP-43 and its derivatives can be easily measured and analyzed in platelet-lysate during the disease prognoses. Intracellular TDP-43 species such as aggregates, truncated TDP-43 fragments, and post-translationally modified TDP-43 have been found in neurodegenerative diseases [53]. The characteristics of post-translationally modified TDP-43 may affect the pathological course of the diseases much earlier than previously thought [54]. There are several supporting studies in the literature cited in the body of this manuscript that have reported TDP-43 levels in serum and brain samples obtained from AD patients. All of these studies provide considerable supporting evidence that a notable percent of AD cases are linked to altered TDP-43. However, the peripheral measurement of TDP-43 and its phosphorylated derivatives are relatively new. In this preliminary study, we have tested our overarching hypothesis stating that peripheral TDP-43 profile may mirror that of brain tissue so that the early diagnosis of AD would be possible before the clinical manifestations are revealed. Peripheral TDP-43 measurement in AD was reported [23]; however, the ELISA method was employed for measuring total TDP-43 in AD patient’s serum. Serum contains some very abundant biomolecules such as albumin and immunoglobulins. These biomolecules may mask the levels of TDP-43 in serum-based assays so that positive recognition of TDP-43 by its selective antibody may be greatly reduced. That is why we have justified utility of anuclear platelet as a biological milieu, which will reflect a more concentrated and encapsulated population of TDP-43 with minimal or no interference of serum albumin, immunoglobulins, and nuclear TDP-43 contribution. We thought that a cell-based TDP-43 chemical modification and aggregation model may be a good strategy [55] to investigate whether peripheral cells would be considered as a platform where the surrogate biomarker such as TDP-43 can be analyzed. Therefore, we have hypothesized that platelet TDP-43 and its phosphorylated derivatives may be considered as a viable surrogate dynamic biomarker.

To identify an AD-selective antibody was the major undertaking in this study. Eight antibodies raised against different regions of TDP-43 as well as phosphorylated derivatives of TDP-43 were tested. They were purchased from different vendors (ProteinTech, Cosmobio-USA, Abcam). We have identified an anti-phospho (S409/410-2) TDP-43 antibody purchased from ProteinTech (Cat#22309-1AP) as an AD-discriminating antibody for AD-platelet lysate phosphorylated TDP-43 content. This antibody did not well-recognize platelet lysate phosphorylated TDP-43 in another neurodegenerative disease, ALS. We have used ALS patient platelet lysate as a negative disease control for testing the selectivity of the antibody that recognized the high levels of pTDP-43 in AD platelet lysate. This observation was unexpected and requires more in depth analysis in larger patient cohort. The alternative explanation would be that AD-selective antibody-producing clones would differ in recognizing the specific modification in TDP-43 in AD platelet lysate.

TDP-43 proteinopathy is characterized by decreased solubility, hyper-phosphorylation and the generation of a 25 kDa C-terminal fragment [16, 56-58]. This section of TDP-43 protein was identified as the disordered region (Fig.1B) which is also a target for a caspase enzyme cleavage [59]. We also have observed ∼35 and ∼25 kDa TDP-43 fragments in platelet lysates obtained from AD patients in early stages of this work (data not shown); we thought that they could represent the degradation products of TDP-43 due to either storage of samples at -80°C for extended period of time or the degradation could be due to the old age of the subjects (Fig. 2C). These fragmented TDP-43 species are more likely encapsulated in immunoreactive inclusion bodies that may be associated TDP-43 relevant disorders [60]. In our view, there are several enzymatic cleavages of TDP-43 that produces cleaved toxic TDP-43 fragments that may be easily phosphorylated. Subsequently, these fragments will first form an aggregation nucleus through protein-protein interactions yielding TDP-43-enriched plaques in CNS tissue. All of these cited studies as well as many others strengthened the concept that TDP-43 protein profile in Alzheimer’s disease may be a good dynamic biomarker that ought to be comprehensively studied. In tissue, cytosolic TDP-43 protein, especially toxic monomers [61], begin to form hyperphosphorylated species. They are sequestered into inclusion bodies as part of the defense mechanism of the organism, suggesting that cytosolic pTDP-43 or detergent-soluble TDP-43 protein is toxic [10]. We neither verified inclusion body presence in platelets nor existing literature reported. What we observed was that cytosolic TDP-43 was present in platelets and phosphorylated species of TDP-43 was elevated in Alzheimer’s disease. We speculate that anuclear platelet cytosol represents the toxic form of TDP-43 species.

How does aberrant brain TDP-43 appear in peripheral blood cells? One explanation might be that the TDP-43 protein has a C-terminus Q/N rich region [62]; therefore, this protein may have the characteristics of prion-like proteins that propagates itself [63, 64] and transfects other cells. Kanouchi et al., have reviewed the recent findings about the prion-like characteristics of TDP-43 propagation and offered the concepts of contiguous and non-contiguous propagation of misfolded proteins including TDP-43 [65]. Considering the leaky BBB in neurodegenerative diseases as well as the ability of platelets to transiently open the BBB via releasing platelet activating factors [37], it is conceivable that aberrant TDP-43 in astrocytes may transfect the blood cells by means of cell-to-cell infection through having access to the blood stream. This concept was recently reviewed on how the aberrant proteins infect the cells [66]. We are well aware that we were unable to obtain the platelets and post-mortem brain tissues from the same subject, which could be the ideal representation of the TDP-43 profile. However, we are searching other AD centers for whether such paired samples would be available for us to verify our preliminary results.

AD-selective anti-phospho (S409/410-2) TDP43 antibody screening could be utilized as part of a diagnostic panel rather than standalone signature marker at this stage. We don’t clearly know the pathobiology of TDP-43 in early stages of AD and other relevant diseases that eventually lead to AD such as mild cognitive impairment (MCI), mild dementia (MD), and frontal lobe dementia (FLT). It would be desirable to obtain platelet samples from patients with AD relevant diseases and to analyze the appearance of phosphorylated TDP-43 in platelets, which help not only monitor the disease prognosis but also contribute to the early diagnosis.

## CONCLUSION AND FUTURE PERSPECTIVE

In this study, we have provided a trend of TDP-43 profiles in AD and age-matched healthy subjects (Fig. 2A, 2C).To our knowledge, we are the first research group to identify the TDP-43 profile in platelets could be considered as a surrogate dynamic biomarker to monitor disease progress as well as the pharmacological treatment response. Our findings about the presence of phosphorylated TDP-43 in platelets from AD patients are intriguing and led us to question as to whether AD is an exclusively CNS or peripheral system disease. This issue is currently questioned and would require some very comprehensive studies [67, 68].

It is a well-known fact that protein aggregations occurs long before the clinical manifestations are revealed [43]. In vitro biophysical studies in cell culture and mouse brain have suggested that TDP-43 naturally tends to form a dimeric protein as cited in a recent review [69]. Could we monitor TDP-43 modifications and aggregations during disease progression? This issue was always a challenge and leads us to plan longitudinal studies in future. Perhaps the platelet TDP-43 approach will make these kinds of studies feasible. As discussed by Budini et al., cell-based TDP-43 aggregation and modifications model is a powerful tool [55] to test novel therapeutic strategies aimed at preventing and/or reducing TDP-43 aggregation in AD.

In the near future, research teams may consider some therapeutic approaches by which cell permeable chemical chaperons that bind to misfolded protein and stabilize the folded state reduce protein misfolding [70]. In normal circumstances, the molecular chaperons and other housekeeping mechanisms ensure that potentially toxic aberrant proteins or pre-fibrillary aggregates are neutralized before they can do cellular damage [71, 72]. Therefore, the researchers need to know the folding features of protein of interest. If we know the folding features of TDP-43 and can measure the occurrence of misfolded, disease prone TDP-43 early enough, we may be able to stabilize the misfolded protein by potential chemical chaperons, which may open up new therapeutical venues for neurodegenerative disease treatment.

Finally, our results suggest that peripheral blood-derived platelets could be used a venue to identify AD-relevant biomarker candidate proteins. We have identified an AD-selective anti-phospho (S409/410-2) TDP-43 Ab in platelet lysates of AD patients. In future studies, we would like to utilize this antibody as a screening tool in large AD patient population and in addition to other AD-relevant diseases such as MCI, MD, and FLT to validate that phosphorylated TDP-43 could be a reliable biomarker candidate. The challenge is to identify a reliable and validated blood-based biomarker(s) during next 4-6 years.

## EXECUTIVE SUMMARY

- TDP-43 and its phosphorylated derivative can be measured in platelet lysate
- A-Phospho (S409/410-2) TDP-43 is identified as a selective antibody for AD patients that discriminates AD from non-demented control and ALS based on platelet analysis of phospho-TDP43.
- This AD-selective antibody may be utilized as a screening tool to strengthen AD diagnosis along with cognitive tests.
- Patient populations with MCI, MD, and FLT need to be assessed for the phosphorylated TDP-43 profile.

## Acknowledgements

AA acknowledges the contributions of the student research fellows of College of Osteopathic Medicine. We are grateful for Drs. Eric Vidoni and Kathy Newman, and KU Medical Center biorepository facilities for providing brain tissues and platelet lysates. We are thankful for Emre Agbas for editing process of this manuscript.

Papers of special note have been highlighted as: (*) of interest; (* *) of considerable interest

## REFERENCES

1 Alzheimer’s disease statistics. http://www.alzheimers.net/resources/alzheimers-statistics/.

2 Henriksen K, O’bryant SE, Hampel H et al. The future of blood-based biomarkers for Alzheimer’s disease. Alzheimers Dement 10(1), 115–131 (2014).

3 Schneider P, Hampel H, Buerger K. Biological marker candidates of Alzheimer’s disease in blood, plasma, and serum. CNS Neurosci. Ther. 15(4), 358–374 (2009).

4 O’bryant SE, Mielke MM, Rissman RA et al. Blood-based biomarkers in Alzheimer disease: Current state of the science and a novel collaborative paradigm for advancing from discovery to clinic. Alzheimers Dement 13(1), 45–58 (2017). (**) This comprehensive article reviews the recent literature on blood-based biomarkers in AD to provide a current state of art.

5 Ritter A, Cummings J. Fluid Biomarkers in Clinical Trials of Alzheimer’s Disease Therapeutics. Front. Neurol. 6 186 (2015). (*) This paper reviews the literature regarding the existing and emerging fluid biomarkers and to examine how fluid biomarkers have been incorporated into clinical trials.

6 Hertze J, Minthon L, Zetterberg H, Vanmechelen E, Blennow K, Hansson O. Evaluation of CSF biomarkers as predictors of Alzheimer’s disease: a clinical follow-up study of 4.7 years. J. Alzheimers Dis. 21(4), 1119–1128 (2010).

7 Rosa-Neto P, Hsiung GY, Masellis M, Participants C. Fluid biomarkers for diagnosing dementia: rationale and the Canadian Consensus on Diagnosis and Treatment of Dementia recommendations for Canadian physicians. Alzheimers Res. Ther. 5(Suppl 1), S8 (2013).

8 Mattsson N, Zetterberg H, Janelidze S et al. Plasma tau in Alzheimer disease. Neurology 87(17), 1827–1835 (2016).

9 Lovheim H, Elgh F, Johansson A et al. Plasma concentrations of free amyloid beta cannot predict the development of Alzheimer’s disease. Alzheimers Dement 13(7), 778–782 (2017).

10 Ugras SE, Shorter J. RNA-Binding Proteins in Amyotrophic Lateral Sclerosis and Neurodegeneration. Neurol. Res. Int. 2012 432780 (2012).

11 Amador-Ortiz C, Lin WL, Ahmed Z et al. TDP-43 immunoreactivity in hippocampal sclerosis and Alzheimer’s disease. Ann. Neurol. 61(5), 435–445 (2007).

12 Baloh RH. TDP-43: the relationship between protein aggregation and neurodegeneration in amyotrophic lateral sclerosis and frontotemporal lobar degeneration. FEBS J. 278(19), 3539–3549 (2011).

13 Buratti E, Baralle FE. The molecular links between TDP-43 dysfunction and neurodegeneration. Adv. Genet. 66 1-34 (2009).

14 Guo W, Chen Y, Zhou X et al. An ALS-associated mutation affecting TDP-43 enhances protein aggregation, fibril formation and neurotoxicity. Nat. Struct. Mol. Biol. 18(7), 822–830 (2011).

15 Geser F, Stein B, Partain M et al. Motor neuron disease clinically limited to the lower motor neuron is a diffuse TDP-43 proteinopathy. Acta Neuropathol. 121(4), 509–517 (2011).

16 Neumann M, Sampathu DM, Kwong LK et al. Ubiquitinated TDP-43 in frontotemporal lobar degeneration and amyotrophic lateral sclerosis. Science 314(5796), 130–133 (2006).

17 Buratti E, Baralle FE. TDP-43: gumming up neurons through protein-protein and protein-RNA interactions. Trends Biochem. Sci. 37(6), 237–247 (2012).

18 Fallini C, Bassell GJ, Rossoll W. The ALS disease protein TDP-43 is actively transported in motor neuron axons and regulates axon outgrowth. Hum. Mol. Genet. 21(16), 3703–3718 (2012).

19 Ayala YM, Zago P, D’ambrogio A et al. Structural determinants of the cellular localization and shuttling of TDP-43. J. Cell Sci. 121(Pt 22), 3778–3785 (2008).

20 Fiesel FC, Kahle PJ. TDP-43 and FUS/TLS: cellular functions and implications for neurodegeneration. FEBS J. 278(19), 3550–3568 (2011).

21 Freeman SH, Spires-Jones T, Hyman BT, Growdon JH, Frosch MP. TAR-DNA binding protein 43 in Pick disease. J. Neuropathol. Exp. Neurol. 67(1), 62–67 (2008).

22 Wilson AC, Dugger BN, Dickson DW, Wang DS. TDP-43 in aging and Alzheimer’s disease -a review. Int. J. Clin. Exp. Pathol. 4(2), 147–155 (2011).

23 Foulds P, Mcauley E, Gibbons L et al. TDP-43 protein in plasma may index TDP-43 brain pathology in Alzheimer’s disease and frontotemporal lobar degeneration. Acta Neuropathol. 116(2), 141–146 (2008).

24 Verstraete E, Kuiperij HB, Van Blitterswijk MM et al. TDP-43 plasma levels are higher in amyotrophic lateral sclerosis. Amyotroph. Lateral. Scler. 13(5), 446–451 (2012).

25 Yamashita T, Teramoto S, Kwak S. Phosphorylated TDP-43 becomes resistant to cleavage by calpain: A regulatory role for phosphorylation in TDP-43 pathology of ALS/FTLD. Neurosci. Res. 107 63–69 (2016).

26 Uchida A, Sasaguri H, Kimura N et al. Non-human primate model of amyotrophic lateral sclerosis with cytoplasmic mislocalization of TDP-43. Brain 135(Pt 3), 833–846 (2012).

27 Italiano Jr. JE, Hartwig JH. Megakaryocyte and platelet structure. In: Hematology, Basic Principles and Practice, Hoffman R, Benz J, E.J., Shattil SJ et al. (Ed.(Eds). Elsevier USA 1873–1880 (2005).

28 Stahl SM, Meltzer HY. A kinetic and pharmacologic analysis of 5-hydroxytryptamine transport by human platelets and platelet storage granules: comparison with central serotonergic neurons. J. Pharmacol. Exp. Ther. 205(1), 118–132 (1978).

29 Veitinger M, Varga B, Guterres SB, Zellner M. Platelets, a reliable source for peripheral Alzheimer’s disease biomarkers? Acta Neuropathol Commun 2 65 (2014).

30 Joseph R, Tsering C, Grunfeld S, Welch KM. Serotonin may have neurotoxic properties. Neurosci. Lett. 136(1), 15–18 (1992).

31 Fisar Z, Hroudova J, Hansikova H et al. Mitochondrial Respiration in the Platelets of Patients with Alzheimer’s Disease. Current Alzheimer research 13(8), 930–941 (2016).

32 Inekci D, Jonesco DS, Kennard S, Karsdal MA, Henriksen K. The potential of pathological protein fragmentation in blood-based biomarker development for dementia -with emphasis on Alzheimer’s disease. Front. Neurol. 6 90 (2015).

33 Johnston JA, Liu WW, Coulson DT et al. Platelet beta-secretase activity is increased in Alzheimer’s disease. Neurobiol. Aging 29(5), 661–668 (2008).

34 Youmans KL, Wolozin B. TDP-43: a new player on the AD field? Exp. Neurol. 237(1), 90–95 (2012). (**) This article reviewes TDP-43 as a new player in Alzheimer’s disease.

35 Marksteiner J, Humpel C. Platelet-derived secreted amyloid-precursor protein-beta as a marker for diagnosing Alzheimer’s disease. Curr. Neurovasc. Res. 10(4), 297–303 (2013).

36 Junt T, Schulze H, Chen Z et al. Dynamic visualization of thrombopoiesis within bone marrow. Science 317(5845), 1767–1770 (2007).

37 Fang W, Zhang R, Sha L et al. Platelet activating factor induces transient blood-brain barrier opening to facilitate edaravone penetration into the brain. J. Neurochem. 128(5), 662–671 (2014).

38 Qureshi AH, Chaoji V, Maiguel D et al. Proteomic and phospho-proteomic profile of human platelets in basal, resting state: insights into integrin signaling. PLoS One 4(10), e7627 (2009).

39 Morris JC, Weintraub S, Chui HC et al. The Uniform Data Set (UDS): clinical and cognitive variables and descriptive data from Alzheimer Disease Centers. Alzheimer Dis. Assoc. Disord. 20(4), 210–216 (2006).

40 BCA protein assay method. http://www.assay-protocol.com/biochemistry/BCA-assay.

41 Jacks A, Babon J, Kelly G et al. Structure of the C-terminal domain of human La protein reveals a novel RNA recognition motif coupled to a helical nuclear retention element. Structure 11(7), 833–843 (2003).

42 Piovesan D, Tabaro F, Micetic I et al. DisProt 7.0: a major update of the database of disordered proteins. Nucleic Acids Res. 45(D1), D219–D227 (2017).

43 Ross CA, Poirier MA. Protein aggregation and neurodegenerative disease. Nat. Med. 10 Suppl S10-17 (2004).

44 Cascella R, Capitini C, Fani G, Dobson CM, Cecchi C, Chiti F. Quantification of the Relative Contributions of Loss-of-function and Gain-of-function Mechanisms in TAR DNA-binding Protein 43 (TDP-43) Proteinopathies. J. Biol. Chem. 291(37), 19437–19448 (2016).

45 Bozzo F, Salvatori I, Iacovelli F et al. Structural insights into the multi-determinant aggregation of TDP-43 in motor neuron-like cells. Neurobiol. Dis. 94 63–72 (2016).

46 Jack CR, Jr., Knopman DS, Jagust WJ et al. Hypothetical model of dynamic biomarkers of the Alzheimer’s pathological cascade. Lancet Neurol. 9(1), 119–128 (2010).

47 Kim J, Choi IY, Duff KE, Lee P. Progressive Pathological Changes in Neurochemical Profile of the Hippocampus and Early Changes in the Olfactory Bulbs of Tau Transgenic Mice (rTg4510). Neurochem. Res. 42(6), 1649–1660 (2017).

48 Adav SS, Sze SK. Insight of brain degenerative protein modifications in the pathology of neurodegeneration and dementia by proteomic profiling. Mol. Brain 9(1), 92 (2016).

49 Kadokura A, Yamazaki T, Lemere CA, Takatama M, Okamoto K. Regional distribution of TDP-43 inclusions in Alzheimer disease (AD) brains: their relation to AD common pathology. Neuropathology 29(5), 566–573 (2009). (*) This paper discusses abnormal TDP-43 deposition and AD pathology may be independently occur.

50 Uryu K, Nakashima-Yasuda H, Forman MS et al. Concomitant TAR-DNA-binding protein 43 pathology is present in Alzheimer disease and corticobasal degeneration but not in other tauopathies. J. Neuropathol. Exp. Neurol. 67(6), 555–564 (2008).

51 Josephs KA, Whitwell JL, Knopman DS et al. Abnormal TDP-43 immunoreactivity in AD modifies clinicopathologic and radiologic phenotype. Neurology 70(19 Pt 2), 1850–1857 (2008).

52 Bao X, Pal R, Hascup KN et al. Transgenic expression of Glud1 (glutamate dehydrogenase 1) in neurons: in vivo model of enhanced glutamate release, altered synaptic plasticity, and selective neuronal vulnerability. J. Neurosci. 29(44), 13929–13944 (2009).

53 Lagier-Tourenne C, Polymenidou M, Cleveland DW. TDP-43 and FUS/TLS: emerging roles in RNA processing and neurodegeneration. Hum. Mol. Genet. 19(R1), R46–64 (2010).

54 Bowden HA, Dormann D. Altered mRNP granule dynamics in FTLD pathogenesis. J. Neurochem. 138 Suppl 1 112–133 (2016).

55 Budini M, Buratti E, Stuani C et al. Cellular model of TAR DNA-binding protein 43 (TDP-43) aggregation based on its C-terminal Gln/Asn-rich region. J. Biol. Chem. 287(10), 7512–7525 (2012).

56 Arai T, Hasegawa M, Akiyama H et al. TDP-43 is a component of ubiquitin-positive tau-negative inclusions in frontotemporal lobar degeneration and amyotrophic lateral sclerosis. Biochem. Biophys. Res. Commun. 351(3), 602–611 (2006).

57 Hasegawa M, Arai T, Nonaka T et al. Phosphorylated TDP-43 in frontotemporal lobar degeneration and amyotrophic lateral sclerosis. Ann. Neurol. 64(1), 60–70 (2008). (**) This paper identified multiple phosphorylation sites in deposited TDP-43. Must read article

58 Zhang YJ, Xu YF, Cook C et al. Aberrant cleavage of TDP-43 enhances aggregation and cellular toxicity. Proc Natl Acad Sci U S A 106(18), 7607–7612 (2009).

59 Li Q, Yokoshi M, Okada H, Kawahara Y. The cleavage pattern of TDP-43 determines its rate of clearance and cytotoxicity. Nat Commun 6 6183 (2015).

60 Zhang YJ, Xu YF, Dickey CA et al. Progranulin mediates caspase-dependent cleavage of TAR DNA binding protein-43. J. Neurosci. 27(39), 10530–10534 (2007).

61 Wang YT, Kuo PH, Chiang CH et al. The truncated C-terminal RNA recognition motif of TDP-43 protein plays a key role in forming proteinaceous aggregates. J. Biol. Chem. 288(13), 9049–9057 (2013).

62 Fuentealba RA, Udan M, Bell S et al. Interaction with polyglutamine aggregates reveals a Q/N-rich domain in TDP-43. J. Biol. Chem. 285(34), 26304–26314 (2010).

63 Holmes BB, Diamond MI. Cellular mechanisms of protein aggregate propagation. Curr. Opin. Neurol. 25(6), 721–726 (2012).

64 Couthouis J, Hart MP, Shorter J et al. A yeast functional screen predicts new candidate ALS disease genes. Proc Natl Acad Sci U S A 108(52), 20881–20890 (2011).

65 Kanouchi T, Ohkubo T, Yokota T. Can regional spreading of amyotrophic lateral sclerosis motor symptoms be explained by prion-like propagation? J. Neurol. Neurosurg. Psychiatry 83(7), 739–745 (2012).

66 Guo JL, Lee VM. Cell-to-cell transmission of pathogenic proteins in neurodegenerative diseases. Nat. Med. 20(2), 130–138 (2014). (**). This paper provides the possible mechanisms on how misfolded proteins are being transmitted cell-to-cell.

67 Swerdlow RH. Personal communication. (2016).

68 Martin LJ. Mitochondrial pathobiology in Parkinson’s disease and amyotrophic lateral sclerosis. J. Alzheimers Dis. 20 Suppl 2 S335–356 (2010).

69 Sun Y, Chakrabartty A. Phase to Phase with TDP-43. Biochemistry 56(6), 809–823 (2017).

70 Cohen FE, Kelly JW. Therapeutic approaches to protein-misfolding diseases. Nature 426(6968), 905–909 (2003).

71 Hartl FU, Hayer-Hartl M. Molecular chaperones in the cytosol: from nascent chain to folded protein. Science 295(5561), 1852–1858 (2002).

72 Sherman MY, Goldberg AL. Cellular defenses against unfolded proteins: a cell biologist thinks about neurodegenerative diseases. Neuron 29(1), 15–32 (2001).

